# Differential expression of transposable elements and upregulation of nucleic acid sensing pathways in pterygium

**DOI:** 10.1101/2025.01.26.633420

**Authors:** Patrick C. Demkowicz, Emma Hammes, Despoina Theotoka, Jessica Chow, Mathieu F. Bakhoum

## Abstract

**Purpose:** The mechanisms that lead to pterygium pathogenesis are poorly understood. Ultraviolet (UV) exposure is a leading environmental risk factor. We propose that UV triggers the de-repression of transposable elements (TEs), including endogenous retroviruses (ERVs), and subsequently activate double-stranded RNA and DNA sensors such as RIG-I and cGAS.

**Methods:** In the present study, we analyzed publicly available RNA sequencing data from pterygium and healthy conjunctiva specimens. We used Ingenuity Pathway Analysis (IPA) to examine the relative expression of nucleic acid sensing pathways and related processes. We also used Telescope to identify TEs. We then repeated this analysis using RNA collected from cultured human conjunctival fibroblasts exposed over multiple days to a range of UV intensities.

**Results:** We found that pathways involved in cellular stress, nucleic acid sensing, and innate immune signaling were upregulated in both cell lines exposed to increased amounts of UV energy as well as pterygium. We also observed differential expression of TEs.

**Conclusions:** Taken together, our findings suggest a mechanistic link between environmental factors and pterygium. Understanding the molecular pathways activated by UV exposure may aid in developing therapeutic strategies.

## INTRODUCTION

Pterygium is common ocular surface disorder characterized by a hyperplastic, fibrovascular growth that extends from the conjunctiva onto the cornea.^1, 2^ Worldwide, the overall prevalence of pterygium is about 12% (95% CI 11-14%).^3–5^ Although benign, its encroachment on the corneal surface can significantly impair vision through induction of astigmatism and corneal scarring.^6–8^

The pathophysiological mechanisms underlying pterygium formation remain poorly understood. Several factors have been implicated in its pathogenesis including viral infections, oxidative stress and genetic predisposition.^5, 9–12^ Among established risk factors, sunlight exposure is the most significant one.^3–5, 13–16^ In fact, pterygium is more common in areas with high sun exposure.^17^ Inflammation has also been implicated in pterygium pathogenesis, and viral triggers have been suggested to play a central role.^10, 12, 18^ However, studies have generally failed to identify pathogens such as human papillomavirus (HPV) or herpes simplex virus (HSV) as causative agents in pterygium.^19–25^

In exploring the potential link between UV exposure and inflammation, we hypothesized that chronic UV exposure might induce an inflammatory response through de-repression of transposable elements (TEs), including human endogenous retroviruses (HERVs) and long interspersed elements (LINEs). These are typically silenced by host regulatory mechanisms, and their transcriptional activation could result in the production of RNA and DNA that act as pathogen-associated molecular patterns (PAMPs), thereby stimulating innate inflammatory responses through pattern recognition receptors (PRRs) such as RIG-I/MDA5/MAVS and the cGAS-STING pathways.^26, 27^

To test this hypothesis, we analyzed transcriptional profiles of pterygium and healthy conjunctiva specimens, and assessed the transcriptional and functional responses of primary and immortalized human conjunctival fibroblasts to UV exposure. Our findings demonstrate evidence of de-repression of TEs in pterygium specimens relative to healthy conjunctiva, and in conjunctival fibroblasts upon UV exposure. This was accompanied by upregulation of genes implicated in cell-intrinsic inflammation. These results suggest a mechanistic link between two hallmarks of pterygium – UV exposure and inflammation – and provide a foundation for future therapeutic strategies targeting these pathways.

## METHODS

### Public Datasets

Massive Analysis of cDNA Ends (MACE) RNA sequencing reads from 8 human pterygium samples and 8 healthy human conjunctiva samples were downloaded from Sequence Read Archive (SRA) (accession PRJNA655558) in the form of raw fastq files.^28^ Each condition contained specimens from both male and female subjects; the metadata provided with the sequences did not include sex as a variable. As a result, our study did not consider sex as a biological variable.

### Ultraviolet Radiation Generation

A Stratagene Stratalinker UV 1800 Crosslinker was fitted with one USHIO G8T5E 7.2W Midrange UV-B 306nm Blacklight Tube. To determine the amount of time needed to achieve specific cumulative doses of UV radiation (J/m^2^), a UV light meter was used to measure the average intensity (uW/cm^2^) emitted within prespecified intervals (2 s, 5 s, 10 s) across varying lengths of time (40 s, 100 s, and 200 s, respectively). This was repeated three times for each pair of interval/duration settings. Calibration curves plotting time versus cumulative radiation were calculated from these data and linear regression was performed (Supp. Fig. 5). We used the briefest (with narrowest measurement intervals) calibration curves to interpolate. We estimated that 30 s achieved 100 J/m^2^, 48 s achieved 200 J/m^2^, and 113 s achieved 500 J/m^2^.

### Cell Culture

#### Immortalized Human Conjunctival Fibroblasts - Short-Term UV Exposure

For downstream RNA sequencing, Immortalized human conjunctival fibroblasts (IHCFs; ABM, Inc., Cat. T0375) were plated in Dulbecco’s Modified Eagle Medium (DMEM) with 10% fetal bovine serum and 1% penicillin/streptomycin at a density of 10,000 cells/cm^2^ in two poly-l-lysine (PLL)-coated 6-well plates (Corning) (“Day 0”). Each row of three wells was assigned to a different condition (0 J/m^2^, 100 J/m^2^, 200 J/m^2^, and 500 J/m^2^). The day after plating (Day 1), media was removed and replaced with phosphate-buffered saline (PBS). With all other conditions covered with aluminum foil, the plate was placed with its cover on in the crosslinker machine and exposed to UV-B radiation for a predetermined amount of time. The PBS was then replaced with fresh media and the plate was returned to the incubator. This procedure was repeated for a total of five days for the RNA-sequencing experiment. RNA was harvested on Day 6.

#### Primary Human Conjunctival Fibroblasts - Short-Term UV Exposure

For downstream RNA sequencing, primary human conjunctival fibroblasts (PHCFs; Sciencell, Inc., Cat. 6570) were plated in Sciencell medium at a density of 5000 cells/cm^2^ in two poly-l-lysine (PLL)-coated 6-well plates (Corning) (“Day 0”). Each row of three wells was assigned to a different condition (0 J/m^2^, 100 J/m^2^, 200 J/m^2^, and 500 J/m^2^); the 100 J/m^2^ condition was not sent for sequencing based on preliminary analysis of data from short-term exposed IHCFs. PHCFs were exposed for a total of five days for the RNA-sequencing experiment. RNA was harvested on day 6.

### RNA Sequencing

On day six, we extracted total RNA (Qiagen RNeasy kit). The Kapa Hyper prep kit with RiboErase was used to prepare 150 bp (base pair), paired-sequence libraries consisting of both coding and non-coding RNA. Each sample was sequenced on two lanes of a NovaSeq (Illumina) to a total read depth of 100 million. Raw, zipped fastq files were obtained for downstream pre-processing.

### RNA Sequence Data Processing

Reads were adapter- and quality-trimmed using Trim Galore with standard settings; the *--paired* flag was used for in-house sequence data. For downstream analysis of gene expression, trimmed reads were mapped to the indexed (generated using an annotation file) human genome (GRCh38, Ensembl) using STAR (version 2.7.7a). The mapped reads were exported as coordinate-sorted BAM files. In addition, the mapping step was run with the *--quantMode GeneCounts* flag to obtain raw read counts for each gene. For the pterygium dataset, counts for the 1st read strand aligned with RNA were exported based upon the results of infer_experiment.py from the RSeQC package (v. 5.0.1 from Bioconda). For in-house dataset, counts for the 2^nd^ read strand aligned with RNA were exported consistent with the library preparation method utilized.

For downstream analysis of TE transcription, reads were also mapped using Bowtie 2 with settings recommended by Bendall et al. *(--very-sensitive-local -k 100 --score-min L,0,1.6*).^29^ The resulting unsorted SAM files were converted to BAM formats and sorted by transcript name using SAMTools.

### Transposable Element Identification

The Telescope program (GitHub v1.0.3) was used to assign reads to HERV and LINE-1 TEs in the retro.hg38.v1 GTF annotation file (Github/telescope_annotation_db). The *telescope assign* command was run with *--max_iter 200, --theta_prior 200000*, and *–stranded_mode* (configured according to the library preparation) flags. Final counts were exported for downstream analysis. Counts per pillion (CPM) were calculated by dividing counts by the number of millions of mapped fragments from Bowtie for a given sample.

### Statistics: Differential Gene Expression Analysis

Raw counts were pre-filtered removing any genes with no counts in any samples. The raw, untransformed counts from the public dataset were then supplied to DESeq2 (R v1.42.1) for differential expression analysis.^30^ We considered genes with absolute log2 fold-change (log2FC) values > |1| for public dataset and P value < 10E-6 as differentially expressed among two conditions.

To identify genes whose expression was significantly correlated (both positively and negatively) with increasing UV dosage, we used the limma package (R v3.58.1) to construct linear models with UV dosage (kJ/m^2^ divided by 100) as the predictor variable.^31^ We used the *treat* function to calculate robust empirical Bayes statistics assuming a minimum log2-fold-change < 1.2 for scientifically meaningful changes and allowing an intensity-dependent trend for the prior variance (*trend = TRUE*).

### Statistics: Principal Components and Hierarchical Clustering

We converted filtered, raw gene counts to log2CPM values, normalized to total gene counts, using EdgeR (R v4.0.16). To normalize TE counts, we divided by millions of reads mapped by Bowtie2 in a given sample. We then took the log2 transform of the normalized counts plus 0.001. Given low overall levels of TE expression, in the public dataset we filtered out TEs with log2CPM < 5 in all samples; a threshold of 1 was used for our analysis of in-house datasets given greater depth of sequencing. We used the *pca* function in PCATools (R v2.14), which removes the bottom 10% least variable genes/TEs, to calculate principal components and then used the *biplot* function to plot the first two principal components. We then plotted the scaled, log2(CPM) values for TEs in heatmaps using the *pheatmap* (R v1.0.12) package and performed hierarchical complete-linkage clustering of both samples and TEs using the correlation distance metric.

### Statistics: Overrepresentation Analysis

Differentially expressed pathways were identified using Ingenuity Pathway Analysis (IPA) software (QIAGEN Inc., https://digitalinsights.qiagen.com/IPA).^42^ Thresholds for selecting analysis gene sets were selected to provide IPA a number of genes within their recommended range. For the public dataset, genes identified in differential expression analysis as having Log2 fold-change > |2| and adjusted P-value < 0.05 were selected for analysis. For the limma trend analysis of IHCFs, genes with a Log2 fold-change > |1| and unadjusted P value < 0.0025 were selected. For the limma trend analysis of PHCFs, genes with a Log2 fold-change > |1| and unadjusted P value < 0.0050. Pathways with -logP > 1.301 were selected as differentially represented and the z-score estimate of relative expression plotted.

We used IPA’s comparison analysis function to produce a matrix of z-scores for pathways enriched across datasets: z-score > 1 and P-value < 0.05. We filtered to the following *reactome* pathways: DNA Repair, Immune system > MHC 1 and MHC II, Immune system > Cytokine signaling > Interferon signaling, Immune system > Cytokine signaling > TNFR2 non-canonical NF-kB signaling, and Innate immune system. We also selected the signaling pathway for UVA/B/C-induced MAPK signaling.

### Gene Set Enrichment Analysis

We used Gene Set Enrichment Analysis (GSEA) software (Broad Institute, https://www.gsea-msigdb.org) to identify Hallmark Human MSigDB gene sets enriched in IHCFs exposed to 500 J/m^2^ for 5 days versus controls (0 J/m^2^).^43^ Log2CPM gene expression values were uploaded to GSEA for this analysis. Gene sets with FDR < 0.25 were selected for plotting.

### Statement on Data Derived from Human Eye Tissues or Cells

All data derived from human clinical specimens were obtained from public sources (SRA accession PRJNA655558).^28^ The human cell lines used in this study were obtained from commercial sources (ABM, Inc., Cat. T0375; ScienCell, Cat. 6570).

### Data Availability

All raw sequencing data generated for this study will be deposited to NCBI’s Gene Expression Omnibus (GEO) database prior to publication.

## RESULTS

First, we analyzed gene expression from eight pterygium and eight healthy conjunctiva specimens.^28^ A total of 2,556 genes were differentially expressed between both conditions, with absolute fold changes (FC) greater than 1 and P-value less than 10E-6 **(Figure 1A)** . Ingenuity Pathway Analysis (IPA) identified 126 pathways that were significantly dysregulated between pterygium and conjunctiva specimens – primarily those involving DNA damage response and nucleic acid sensing pathways **(Fig. 1B; Supp. File 1).** These findings suggest activation of cellular stress responses and innate inflammatory mechanisms in pterygium tissues compared to healthy conjunctiva.

**Figure 1.**
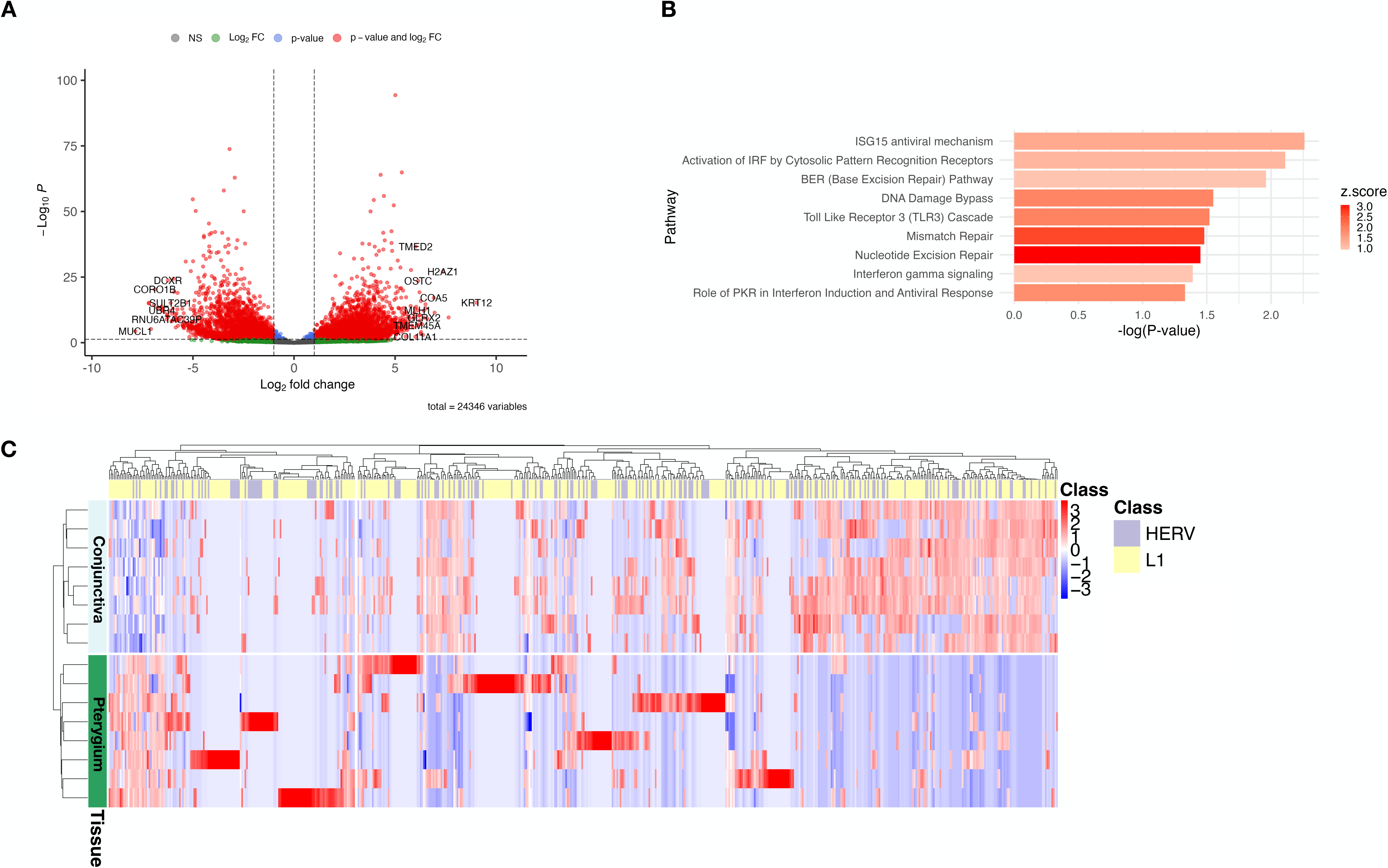
Transcriptional changes in human pterygiumversus conjunctiva. **(A)** Volcano plot showing -log10 of unadjusted P-value from DESeq2 versus the log2 fold change (log2FC). A P-value cutoff of 10E-6 and log2FC > |1| is displayed. A total of 2556 genes were differentially expressed. **(B)** Select DNA damage response, nucleic acid sensing, and downstream inflammatory pathways increased in pterygia vs. healthy conjunctiva as identified in Ingenuity Pathway Analysis (IPA). The x-axis is the -log of the adjusted P-value and the bars are shaded according to the z-score for the pathway comparison (pterygium vs. conjunctiva). Genes were selected for IPA using log2FC > |2| and adjusted P-value < 0.05 thresholds. **(C)** Heatmap and hierarchical clustering of transposable elements including human endogenous retroviruses (HERVs) and LINE-1 elements (L1).

Next, we interrogate for the presence of TEs using Telescope, identifying 592 distinct TEs. Unbiased hierarchical clustering indicated significant differential expression of these TEs between both tissues. Of note, while TE expression in healthy conjunctiva appears homogeneous, TE de-repression in pterygium is stochastic, lacking a conserved TE signature across all eight specimens **(Fig. 1C).**

To establish a causal link between UV exposure and the induction of an inflammatory response, IHCFs were subjected to varying UVB exposures (0, 100, 200, and 500 J/m^2^) over five days. The dosage-dependent impact of UVB on gene expression was evident, with IHCF samples clustering along the first principal component (PC1) by UV dosage **(Fig. 2A).** A total of 3,213 genes were significantly associated with increasing UVB exposure levels (limma-trend analysis; LogFC > |1| and P-value < 0.0025) **(Supp. Fig. 1)**. There were 336 pathways significantly dysregulated secondary to increasing UV dosage, including those involved in DNA damage response, nucleic acid sensing, and downstream inflammatory pathways such as interferon signaling **(Fig. 2B; Supp. File 1).** GSEA of cells exposed to 500 J/m^2^ vs. controls revealed upregulation of inflammatory, UV response, and DNA repair pathways (**Supp. Fig. 2; Supp. File 1**). Furthermore, IHCFs exposed to the highest levels of UVB (500 J/m^2^) exhibited distinct TE expression profiles and clustered separately from those exposed to no or lesser UVB levels **(Fig. 2C).**

**Figure 2.**
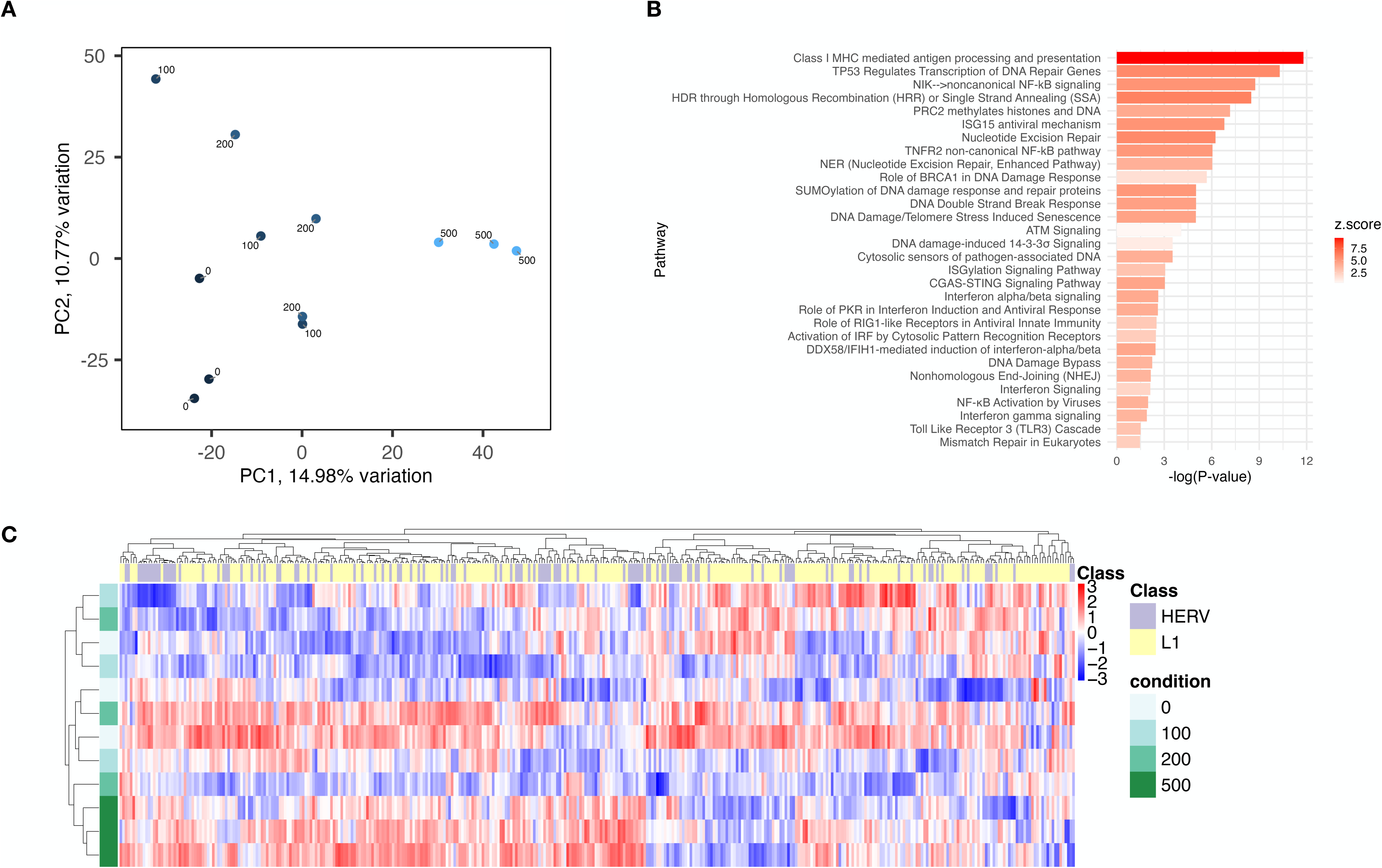
Transcriptional changes in immortalized human conjunctival fibroblasts (IHCFs) exposed to UV. **(A)** Principal component analysis of gene transcription in IHCFs exposed to 0, 200, and 500 J/m^2^ x five days. **(B)** Select DNA damage response, nucleic acid sensing, and downstream inflammatory pathways correlated with increasing UV as identified in Ingenuity Pathway Analysis (IPA). The x-axis is the -log of the adjusted P-value and the bars are shaded according to the z-score. Genes were selected for IPA using Log2 fold-change > |1| and unadjusted P value < 0.0025 thresholds. **(C)** Heatmap and hierarchical clustering of transposable elements.

Similar trends were observed in Primary Human Conjunctival Fibroblast (PHCF) cell lines. PCA of gene transcripts showed clustering by UV dosage **(Supp. Fig. 3A).** A total of 3,306 genes associated with increasing UVB exposure levels (limma-trend analysis; LogFC > |1| and P-value < 0.0050) were selected for IPA **(Supp. Fig. 4)**. IPA identified 418 pathways that were significantly dysregulated upon increasing UV dosage, including pathways involved in DNA damage, nucleic acid sensing, and inflammatory processes **(Supp. Fig. 3B; Supp. File 1).** Lastly, PHCFs exhibited distinct TE expression profiles with respect to UV dosage **(Supp. Fig. 3C).**

Next, we compared genes that were significantly different between pterygium and conjunctiva to genes identified in limma-trend analysis of UV-exposed IHCF and PHCF cells. There were 298 genes (4.4%) common among all gene sets, 194 (2.9%) between pterygium and IHCF gene sets, 1021 (15.0%) between IHCF and PHCF gene sets, and 482 (7.1%) between PHCF and pterygium gene sets **(Fig. 3A).** We also compared core IPA pathways related to DNA repair, MHC signaling, interferon signaling, non-canonical NF-kB signaling, innate immune signaling, and UV response that were identified pterygium, IHCF, and PHCF datasets **(Fig. 3B).** We found substantial concordance with respect to the pathways upregulated in pterygium and positively associated with increasing UV dosage.

**Figure 3.**
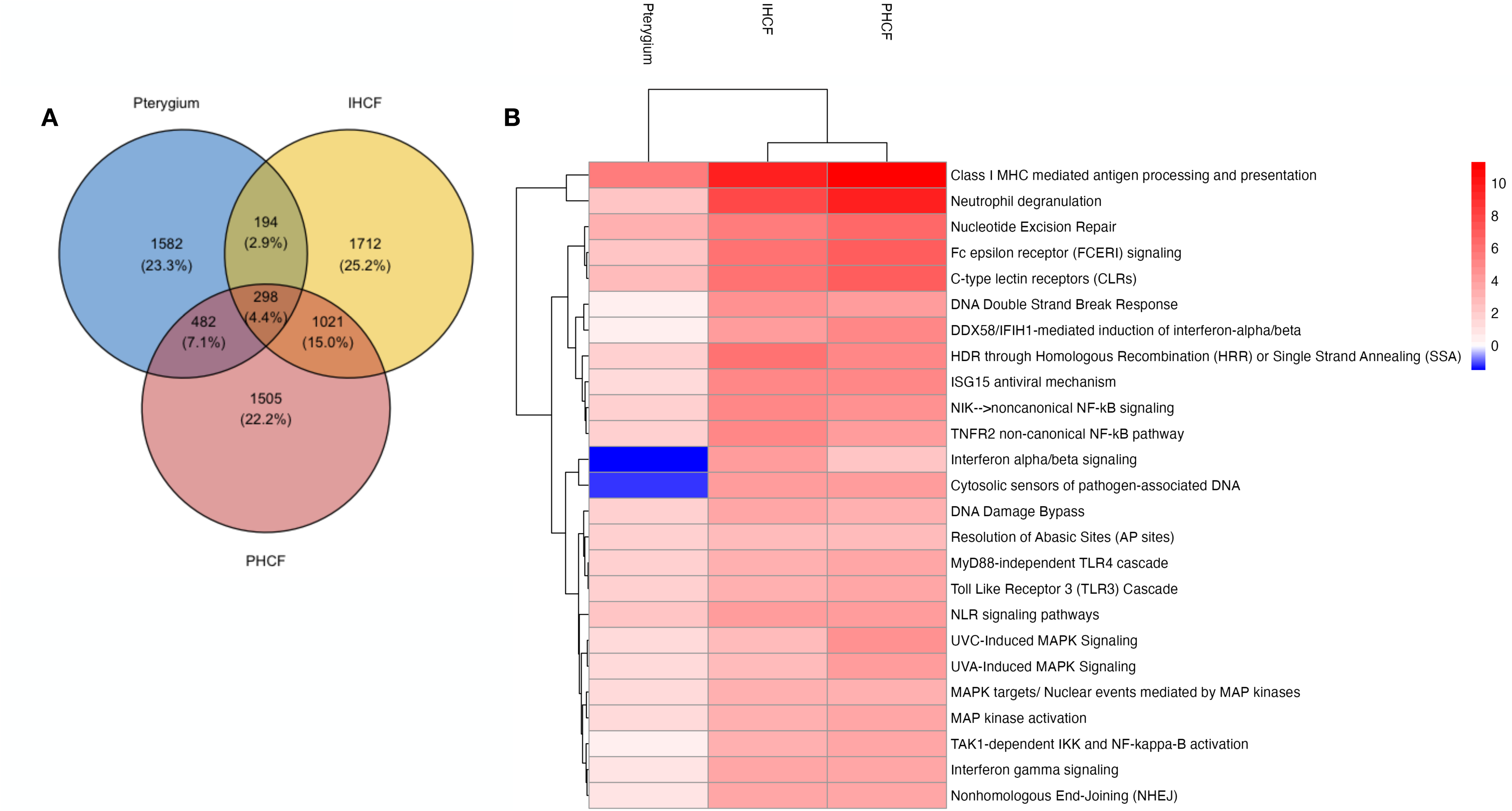
Comparison of human and in vitro datasets. **(A)**Comparison of differentially expressed genes. Thresholds: logFC > |1| for all; P-value < 10E-6 for pterygium, P-value < 0.0025 for immortalized human conjunctival fibroblasts (IHCFs), and P-value < 0.0050 for primary human conjunctival fibroblasts (PHCFs). **(B)** Comparison of select Ingenuity Pathway Analysis pathways. The z-score for each pathway is colored. Pathway threshold: z-score > 1 and P-value < 0.05.

## DISCUSSION

Our study provides compelling evidence for the role of UV exposure in the de-repression of TEs and activation of nucleic acid sensing pathways in pterygium. This suggests a mechanistic link between environmental factors and the pathophysiology of this common ocular surface disorder.

Significant differential expression of TEs was observed in pterygium compared to healthy conjunctiva. While TE expression in conjunctiva was uniform across all samples, their levels varied widely in pterygium specimens, with no conserved TE signature. This stochastic pattern of de-repression highlights the potential role of environmental factors, such as UV radiation, in sporadically triggering TE de-repression rather than a uniform activation across individuals. The differential expression of TEs upon UVB exposure – especially at higher UVB dosages – further establishes a causation between UVB exposure and TE de-repression. Prior studies in skin fibroblasts have similarly noted UVB-induced TE activation.^32^ The mechanisms by which this de-repression occurs are unknown, but possible explanations include UV-induced hypomethylation or H3K27ac redistribution across TE-containing regions.^33, 34^

Presence of inflammatory biomarkers in pterygium has led many to speculate that an infectious agent might trigger its formation, with HPV frequently identified in varying degrees in different studies. However, no definitive proof of this causation has been demonstrated. Our findings suggest an alternative mechanism for the inflammatory response observed in pterygium, independent of an infectious etiology – that is through de-repression of endogenous retroviruses. Transcription of TEs has been proposed to activate innate immune responses via stimulation of pattern recognition receptors (PRRs) such as RIG-I/MDA5/MAVS and cGAS-STING.^26, 27^ This is further supported by our identification of enriched pathways involved in nucleic acid sensing, DNA damage repair, and inflammatory signaling in pterygium, and upon UV exposure. Previous transcriptomic analyses of pterygium specimens have focused on pathways related to changes in biomechanical functions such as cell adhesion and extracellular matrix remodeling.^28, 35–37^ Interestingly, Chen et al. observed that GADD34 – which has been shown to mediate the interferon response to viral dsRNA - comprised a hub in the gene transcription network of a subset of pterygium samples; however, it was found by real-time PCR to be downregulated in pterygium compared to conjunctiva controls.^38–40^

Worldwide, the overall prevalence of pterygium is about 12% (95% CI 11-14%).^3–5^ Individuals with pterygium often complain of discomfort, dry eye symptoms and vision deterioration, secondary to astigmatism.^3, 6–8^ The mainstay treatment for pterygium is surgical excision.^41^ Unfortunately, post-surgical recurrence is not uncommon, occurring in about 7 to 20% of cases even with the use of a conjunctival autograft.^8, 41^ There is a need to develop medical therapy that can be used to stop pterygium progression early in the disease course.^10, 41^ Understanding the molecular pathways activated by UV exposure and their role in pterygium formation can aid in developing preventive and therapeutic strategies. Interventions aimed at modulating the activity of pattern recognition receptors or inhibiting elements of the interferon signaling pathway might help mitigate the progression of pterygium. Further research is needed to delineate the inflammatory contributions of damaged host DNA, TE de-repression, and the overall microenvironment in pterygium pathogenesis.

Several limitations should be considered when interpreting the present study. First, the transcriptional analysis was performed on specimens that were formalin-fixed and obtained at different time points, reducing the integrity of RNA. Second, the technology used to sequence the human pterygium specimens, Massive Analysis of cDNA Ends (MACE), is biased toward poly-adenylated 3’ ends and in this study yielded short single-end reads. Consequently, this limits our ability to detect all TEs, which are typically located in introns or not transcribed into poly-adenylated mRNA. However, for conjunctival fibroblasts we used ribosomal depletion prior to library preparation and sequenced them using paired-end reads. Third, we conducted our *in vitro* experiments using fibroblast-derived cell lines, assuming that these cells are the most important cells implicated in pterygium development. However, the cell type from which pterygia are derived has not been established. Additionally, while our *in vitro* model provides valuable insights, the translation of these findings to in vivo systems and clinical practice requires further validation.

In summary, our findings underscore the critical role of UV-induced TE de-repression and nucleic acid sensing pathway activation in pterygium pathogenesis, offering new avenues for understanding and potentially halting the progression of this common ocular disorder.

## AUTHOR CONTRIBUTIONS

**PD:** Methodology, formal analysis, investigation, writing – original draft, visualization; **EH:** Investigation, writing – review & editing; **DT:** writing – review & editing; **JC:** writing – review & editing; **MB:** Conceptualization, methodology, investigation, resources, writing – original draft, supervision, funding acquisition.

## ACKNOWLEDGEMENTS

This work was supported by NIH Research Grant R21EY035090 from the National Eye Institute. Additionally, MFB is supported by NIH Research Grant P30CA016359 from the National Cancer Institute, by the Office of the Assistant Secretary of Defense for Health Affairs through the Melanoma Research Program under Award No. HT9425-23-1-1070, and a grant from the Connecticut Lions Eye Research Foundation. PD receives fellowship funding from the Yale School of Medicine Richard K. Gershon, M.D. Fund, James G. Hirsch, M.D. Endowed Medical Student Research Fellowship, William U. Gardner Memorial Student Research Fellowship, and Leon Rosenberg, M.D., Medical Student Research Fellowship.

## Notes

**Funding:** This work was supported by NIH Research Grant R21EY035090 from the National Eye Institute.

### Competing Interest Statement

PD owns Roivant Sciences stock options.

