## Supplementary material for "Differential expression of transposable elements and upregulation of nucleic acid sensing pathways in pterygium": Supp. Fig.

### SUPPLEMENTARY FIGURES

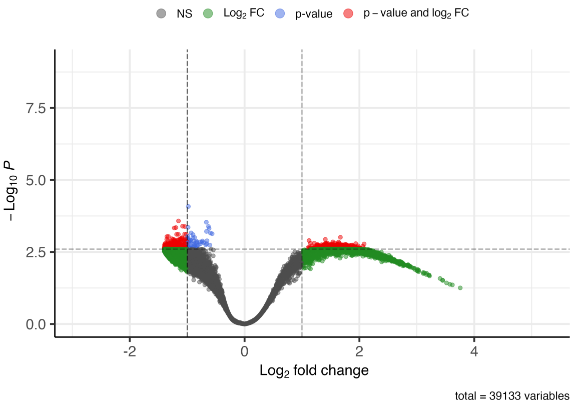


**Supplementary Figure 1. Volcano plot of genes included in limma-trend analysis of immortalized human conjunctival fibroblasts exposed to 0, 100, 200, and 500 J/m$^2$ UV-B x 5 days.** Analysis genes were selected using logFC > |1| and p-value < 0.0025.

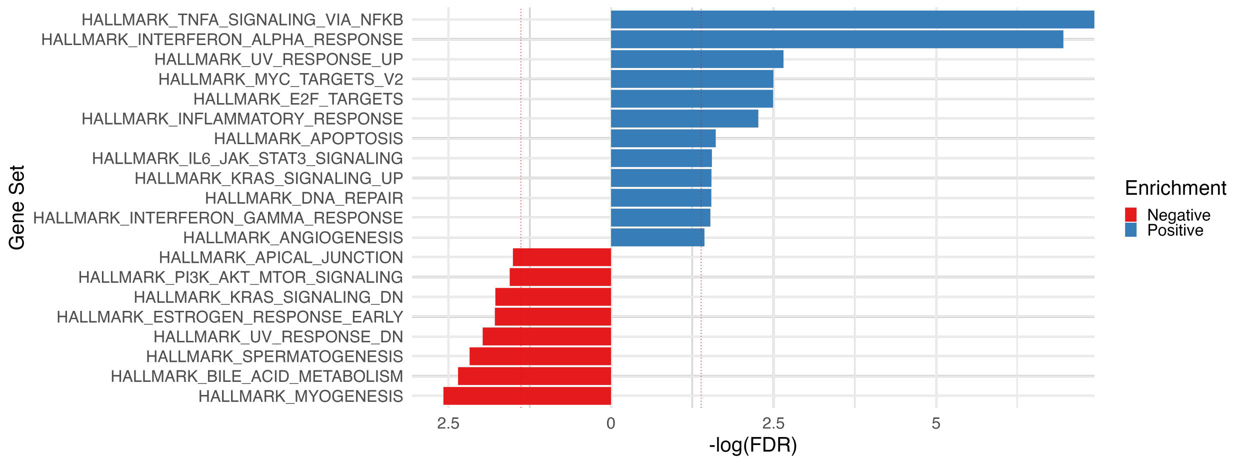


**Supplementary Figure 2. Gene set enrichment analysis results of immortalized human conjunctival cells exposed to 500 J/m$^2$ vs. 0 J/m$^2$ x 5 days.** Gene sets with FDR < 0.25 were selected.

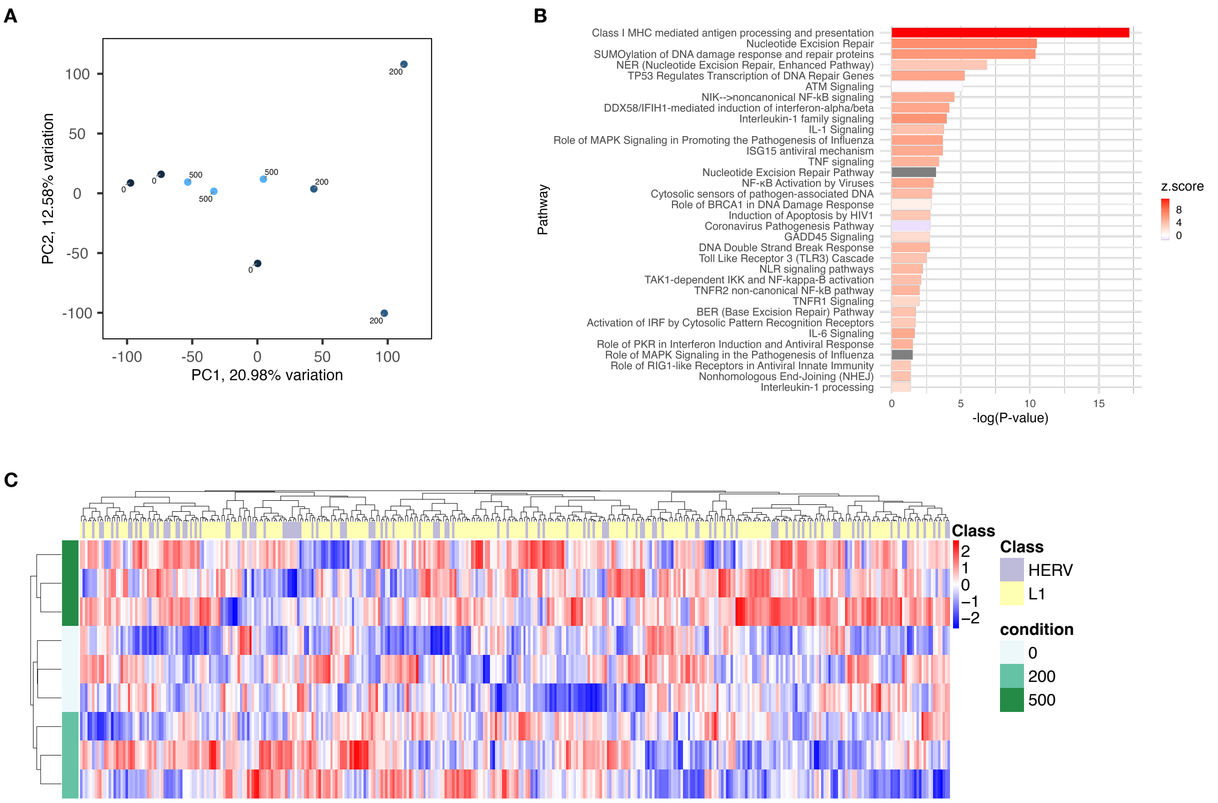


**Supplementary Figure 3. Transcriptional changes in primary human conjunctival fibroblasts (PHCFs) exposed to UV.** (A) Principal component analysis gene transcription in PHCFs exposed to 0, 200, and 500 J/m$^2$ x 5 days. (B) Select DNA damage response, nucleic acid sensing, and downstream inflammatory pathways correlated with increasing UV as identified in Ingenuity Pathway Analysis (IPA). The x-axis is the -log of the adjusted P-value and the bars are shaded according to the z-score. Genes were selected for IPA using Log2 fold-change > |1| and unadjusted P value < 0.005 thresholds. (C) Heatmap and hierarchical clustering of transposable elements.

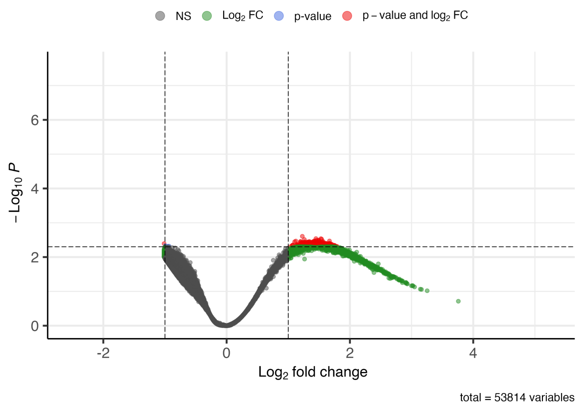


**Supplementary Figure 4. Volcano plot of genes included in limma-trend analysis of primary human conjunctival fibroblasts exposed to 0, 200, and 500 J/m$^2$ x 5 days.** Analysis genes were selected using logFC > |1| and p value < 0.0050.

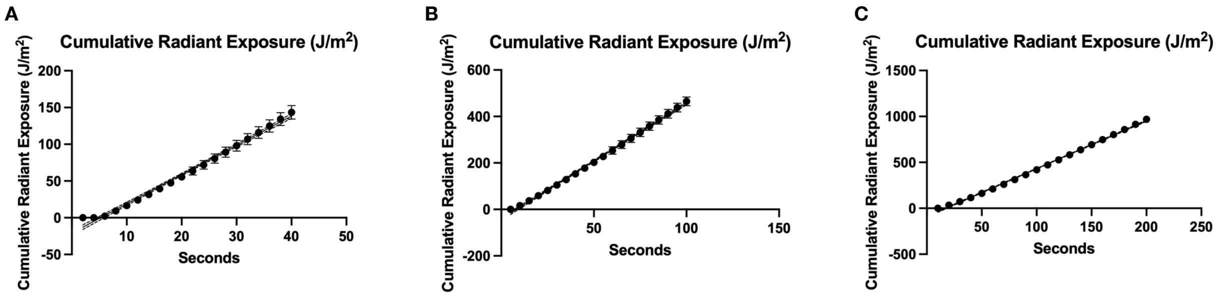


**Supplementary Figure 5. Cumulative radiant exposure (J/m$^2$ of UV-B) vs. time.** Each panel produced using a different probe recording average intensity (uW/cm$^2$) over a different time interval (2 s, 5 s, 10 s). Linear regression performed and used to calculate times needed to administer desired doses of UV-B.
